# Synth4bench: Synthetic Data Generation for Benchmarking Tumor-Only Somatic Variant Calling Algorithms

**DOI:** 10.1101/2024.03.07.582313

**Authors:** Styliani-Christina Fragkouli, Nikos Pechlivanis, Anastasia Anastasiadou, Georgios Karakatsoulis, Aspasia Orfanou, Panagoula Kollia, Andreas Agathangelidis, Fotis Psomopoulos

## Abstract

**Motivation:** Somatic variant calling is a key activity towards identifying genomic alterations; yet, the evaluation of the respective tools remains challenging due to the scarcity of high quality ground truth datasets. To overcome this limitation, we developed synth4bench, a synthetic data generation pipeline for robust benchmarking. Using a systematic process to create distinct synthetic datasets, we thoroughly evaluated five variant callers (Mutect2, FreeBayes, VarDict, VarScan2 and LoFreq). We compared tool outputs against our synthetic ground truth across key sequencing aspects (such as depth and read length) to assess their capacities and shed light on their underlying algorithmic principles.

**Results:** Synth4bench is an approach for evaluating tumor-only somatic variant callers that relies on a systematic definition of fully controlled ground-truth datasets. Our analysis revealed significant inconsistencies among the tool outputs and a strong dependence of caller performance on sequencing parameters. Indels remain the hardest-to-call variant type, driven by errors at low allele frequencies. Algorithmic choice is also critical; the most robust callers displayed the highest precision in allele frequency estimation, while the most sensitive caller was best for maximizing true positive recovery. Conversely, the least suitable caller exhibited systematic errors along with the poorest overall performance. These findings indicate that there isn’t a one-solution-fit-all; sequencing optimization together with caller selection are necessary to maximize sensitivity and reliability. Furthermore, the pronounced inconsistencies suggest that current algorithms are not yet able to capture all mutational mechanisms adequately, with the modeling of the underlying processes remaining an open challenge.

**Availability:** code: https://github.com/sfragkoul/synth4bench/ and data: https://zenodo.org/records/16524193

**Graphical Abstract:** 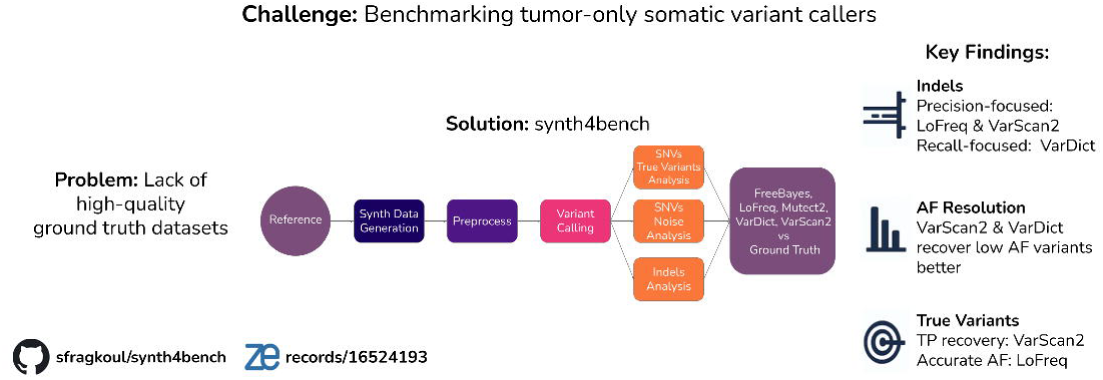

## 1. Introduction: Framing the Challenge

It is commonly known that diseases such as cancer are caused by alterations in the genomic DNA of organisms. These alterations are induced by a plethora of factors such as aging, heredity, lifestyle, viral infections and more. High throughput sequencing (HTS) technologies have greatly assisted in capturing the actual genomic complexity of cancer, through the identification of thousands of somatic mutations related to a variety of human cancers [1,2]. The accurate detection of somatic variants, especially those at low frequencies, plays a pivotal role in unraveling genomic alterations that represent the actual drivers of these complex diseases [3,4].

To date, a large series of bioinformatics tools are available for the detection of all different types of mutations, including single nucleotide variants (SNVs), as well as small insertions and deletions (indels). Yet, the accuracy and consistency of this process is still challenging due to the absence of standards and criteria [4–16]. Furthermore, in the case of variants identified at a low allele frequency (AF) (i.e. ≤10%), robust identification becomes even more demanding, requiring higher levels of sensitivity across analytical workflows [7,17].

This unresolved issue is evident in the limited agreement among the results produced by different variant callers. To this date, several publications have dealt with the benchmarking and comparison of relevant algorithms [18–20]. To improve reliability, variant calling pipelines have been refined with new methods and strategies designed to reduce discordant and inconsistent results. To name a few of these ensemble approaches, consensus methods [21], majority voting [12], as well as methods from the domain of machine learning such as Bayesian inference [22], decision trees [23] and deep learning [24] have been applied.

Furthermore, the assessment of variant calling capacity has long been hindered by the scarcity of high-quality datasets that could serve as ground truth [12,13]. The implementation of such datasets would render the process of benchmarking more efficient and robust[15,25]. A relevant solution concerns the use of synthetic data from the *in-silico* simulation of synthetic genomics, which has evolved over the last 30 years [26] alongside sequencing technologies [8,27].

In this work we present synth4bench, a bioinformatics pipeline to observe the behavior of variant calling algorithms. To achieve this, we generated a set of synthetic next-generation sequencing (NGS) datasets; at its heart lies NEAT simulator (NExt-generation sequencing Analysis Toolkit) [28] that was utilized for datasets generation. Varying values of some intrinsic parameters of NGS data, which concerned the sequencing depth and read length, were used in order to study the behavior of different variant callers and their response to their variation. Our results demonstrate that synth4bench assisted significantly in the process of variant calling, by shedding light on the behavior and decisions of the studied algorithms, as well as in outlining their patterns of their responses to different NGS-related parameters.

## 2. Experimental Design and Methods: Crafting the Pipeline

Our pipeline takes as input a reference human genome, the NGS-related parameters (as set by the user) and outputs the comparison of results between each variant caller and the ground truth. The in-between steps include the preprocessing required before moving to the variant calling process, as well as the downstream analysis of reported variants that is necessary to extract the required information, as depicted in **Figure 1**. The comparison concerns results from different analyses, which were implemented in a way to adequately investigate both SNVs as well as indels. A list of all the steps and used tools can be found in **Table 1**.

**Figure 1:**
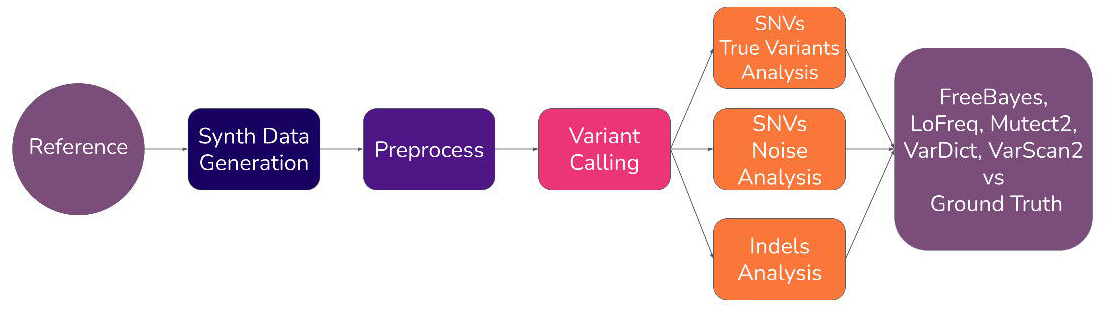
Schematic representation of synth4bench. The circles represent the input and output files, while the rectangles represent the in-between processes. Three different analyses (two for SNVs true variants and noise, and one for indels) follow after the step of variant calling.

**Table 1:** All steps and respective tools, as implemented by synth4bench for the processes of data generation, pre-process, variant calling and benchmarking. Our pipeline can be dynamically adapted; new callers can be added in the future.

| Synthetic Data Generation Steps | Tool |
| --- | --- |
| generation of individual synthetic ground truth files | NEAT |
| merge individual bam ground truth files | samtools merge |
| index individual vcf ground truth files | bcftools index |
| merge individual vcf ground truth files | bcftools merge |
| normalize merged vcf ground truth file | bcftools norm |
| **Common Pre-process Steps** | **Tool** |
| sort merged bam ground truth file | samtools sort |
| clean merged bam ground truth file | samtools view |
| alignment ground truth statistics | samtools flagstat |
| add read group tags in a merged bam ground truth file | samtools addreplacerg |
| compute ground truth coverage per position | samtools depth |
| index all bam ground truth files | samtools index |
| get low-level information about data at each position for all bam ground truth files | bam-readcount |
| **Mutect2 Specific Steps** | **Tool** |
| call Mutect2 to perform variant calling | gatk Mutect2 |
| normalize Mutect2 vcf file | bcftools norm |
| **FreeBayes Specific Steps** | **Tool** |
| call FreeBayes to perform variant calling | FreeBayes |
| Modify header of FreeBayes vcf file | bcftools reheader |
| normalize FreeBayes vcf file | bcftools norm |
| **LoFreq Specific Steps** | **Tool** |
| add indel qualities to ground truth merged bam file | lofreq indelqual |
| call LoFreq to perform variant calling | lofreq call |
| Modify header of LoFreq vcf file | bcftools reheader |
| normalize LoFreq vcf file | bcftools norm |
| **VarScan2 Specific Steps** | **Tool** |
| get "pileup" textual format from ground truth merged bam file | samtools mpileup |
| call VarScan2 to perform variant calling | varscan pileup2cns |
| convert VarScan2 tsv file to vcf file | extra python script |
| normalize VarScan2 vcf file | bcftools norm |
| **VarDict Specific Steps** | **Tool** |
| call VarDict to perform variant calling | vardict |
| convert VarDict txt file to vcf file | var2vcf |
| Modify header of VarDict vcf file | bcftools reheader |
| normalize VarDict vcf file | bcftools norm |

| Benchmarking Steps | Tool |
| --- | --- |
| call ground truth and callers' files to compare | Synth4bench R scripts |

Aligning with open science practices [29], all necessary resources required for the execution of synth4bench are available and ready to use. The source code is available on GitHub (https://github.com/sfragkoul/synth4bench/) along with the appropriate documentation and a user guide for easier implementation. All synthetic datasets are available in Zenodo (https://zenodo.org/records/16524193).

### 2.1 Generate the synthetic ground truth datasets

Due to its central role within our pipeline, the generation of synthetic human genomics data was a task of high importance. The selected simulator was NEAT, due to its adequate precision [27] as well as flexibility, based on its wide list of available parameters compared to other simulating algorithms [30].

A list of the parameters of NEAT and a short description are presented in **Section S1** of the Supplementary Material. The reference human genome used during the data generation was hg38. In regard to the generation of all datasets, the parameter to determine the seed was used to ensure reproducibility. All used seed values were stored in a database along with the parameters for all generated datasets; these can be found in **Section S2** of the Supplementary Material.

One of the main aims of this work was to better understand the behavior of variant callers. To this end, it was crucial to test their response to data with varying values within the genomic feature. For this reason, we constructed a feature space by using ranging values for given characteristics. Coverage, which refers to the count of reads covering each base of the sequenced DNA, is a feature that plays an important role when dealing with NGS-related errors. One approach to ensure accurate base calling at each given position was by increasing the coverage [31]. In order to assess in-depth the behavior of variant callers, the coverage values that were chosen ranged from 300x to 5000x, as shown in **Section S1** of the Supplementary Material. Another chosen feature was the read length of NGS data; lengths ranging from 50 base pairs (bp) to 300bp were put to test. This approach was applied to cover all possible scenarios for a given fragment (in our case with a mean length of 300bp and standard deviation of 30bp). Grid experiments were performed to cover all the constructed feature space and can be found in **Section S4** of the Supplementary Material. Since we worked with a human reference genome, the ploidy parameter was set to 2 and the average mutation rate was fixed to 0.1.

Although some of the aforementioned values are rarely found in wet lab experiments, we decided to explore them nonetheless, since we took advantage of the fast and inexpensive generation that synthetic data offer.

Finally, the *bam-readcount* package [31] was implemented to obtain low-level information for all positions in the ground truth Binary Alignment Map (BAM) files, thereby enhancing the benchmarking process.

### 2.2 Design the benchmarking

Accurately mapping reads to the reference by using alignment tools is important when identifying genetic variants. However, it is known that the choice of a variant caller exerts a greater influence on the detection of variants [15]. Thanks to the fact that NEAT produces aligned reads, we were able to skip the alignment step and focus merely on the variants’ detection, generated directly in the BAM format.

To emphasize low-frequency variants, all BAM and Variant Call Format (VCF) files from independent runs were merged into a single ground truth file, as shown in **Figure 2** and discussed in the relevant section. During the preprocessing phase, some steps were common for all callers, such as indexing. On the other hand, each variant caller has its own demands. All necessary steps, both common and individual, were incorporated in our pipeline using SAMtools and bcftools [32] and can be found in **Section S.1** of the Supplementary Material.

**Figure 2:**
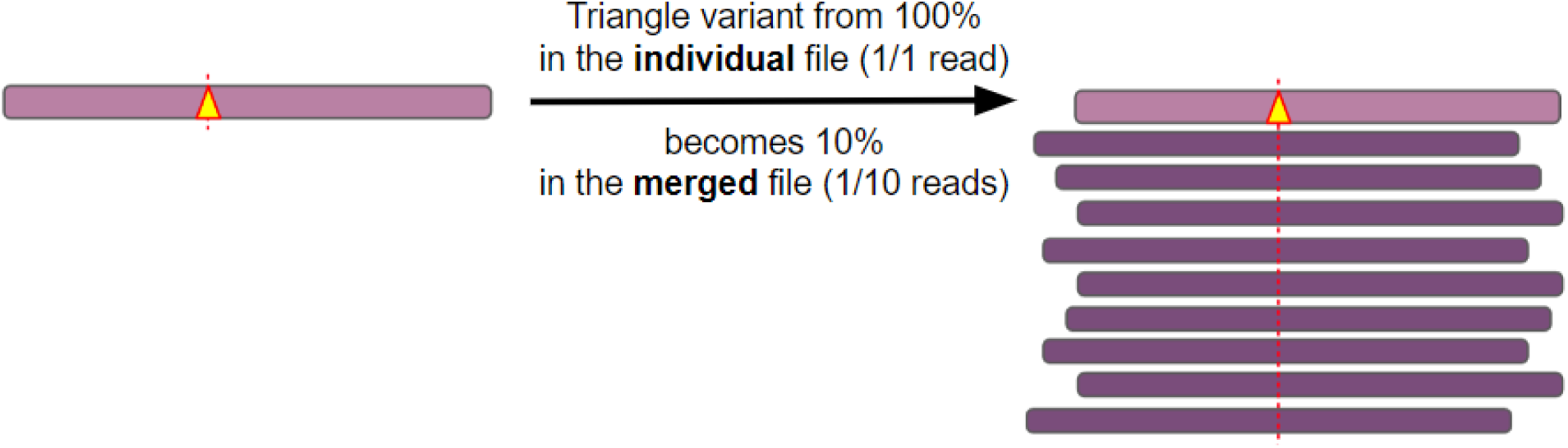
Schematic overview of the approach used to generate low-AF ground truth datasets and define true SNVs. Merging individual read groups led to the recalculation of variant AFs, causing initially high-frequency variants (e.g., 100%) to become “diluted” (e.g., ∼10%) in the combined dataset. Variants unique to the remaining reads (dark purple) were treated as noise.

Our primary goal in this study was to evaluate the technical performance of variant callers. To maintain a consistent and unbiased benchmark, variant annotation was not included and all chromosomal positions were treated equally. This approach focuses on methodological aspects, without considering biological context such as genomic regions or mutation hotspots, which remain important but are beyond the scope of this analysis.

#### 2.2.1 Choosing Variant Callers

As previously noted [15], the nature of variants can largely determine the choice of variant caller. In this study, we focused on popular variant callers, capable of detecting both SNVs and indels [4,6,7,10,11,14–16]. Furthermore, only primary and openly available algorithms were considered. Based on these criteria, the five popular variant callers selected for benchmarking were Mutect2 from GATK 4.3.0.0 [33], FreeBayes 1.3.6 [34], LoFreq 2.1.5 [35], VarScan2 2.4.6 [36] and VarDict 1.8.3 [37], all installed via a conda environment for easier implementation. Ensemble methods were not included in our study. A candidate that was ultimately left out was Strelka2 [38], since it operates only in matched tumor-normal DNA samples. All selected algorithms accept BAM files as input and produce VCF files as output. The sole exception was VarScan2, which required a pile up file as input while additional processing was needed to convert its output from Tab-Separated Values (TSV) format to VCF. All variant callers were run according to the indicated best practices [39–43].

#### 2.2.2 Modelling low-frequency variants

To meet the objectives of our study, we ensured that variant frequencies remained consistently below 10% (AF < 10%) across the three independent analyses. To achieve this, we generated a number *N* of individual datasets with the same NGS characteristics, each of coverage *x*. These individual files were then combined into a single merged file, yielding a final coverage *X* equal to:

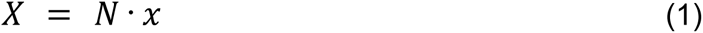

The average AF of variants in each individual file was inversely related to the coverage of the final merged file, as shown in **Equation 2**. If the coverage and AF of a variant in one of the individual files were *x* and *f*, respectively, then the new AF, *F* of the same variant in the merged file would become:

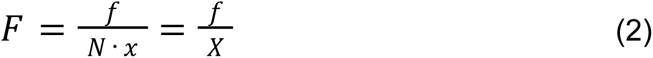

After running a number of experiments to determine the number of individual files *N*, we concluded that the optimal value in our case was 10 because it assisted in reducing sufficiently the overall AFs. Hence, **Equation 1** became *X* = 10*x* and **Equation 2** became 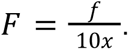

#### 2.2.3 SNVs Independent Analyses of True Variants and Noise

We acknowledged that all data inherently include a true signal that is partially masked by background noise. Based on this and our described methodology, we defined those SNVs that were found in the individual produced files to have AF=100% as “True Variants” and we treated them as the true signal within our data. All other SNVs were treated as noise, despite their AF ranging from small to greater values. The schematic representation of this method is shown in **Figure 2**. The AF of a variant was originally equal to 100% in one of the individual files (i.e. one read with a yellow triangle) and then became 10% in the final merged file (group of a total of 10 reads). In practice, there is usually an overlap among the variants from the individual files; variants from different individual files could be mapped to the same chromosomal position. In the case that the variants happen to be similar, their AF were added. In contrast, if the variants were different, we treated them as background noise.

#### 2.2.4 Indels Independent Analysis

Due to the differences in the nature between SNVs and indels, we deliberately examined them separately. Indels pose a greater challenge, since different variant callers reported the same indels differently, thus hindering the comparison of their calling [44]. To address this challenge, we followed the practice of indel normalization with BCFtools [32]; each variant was normalized based on two conditions; reporting was based on the smallest base position possible, i.e. left-aligned, and the length of each reported variant was the shortest possible, i.e. parsimonious.

As previously discussed [15,16], such complex variants are consistently reported by variant callers. For this reason, indel analysis included some extra steps regarding their normalisation. For example, a deletion that was reported like T>-A was converted to TA>T, while calibration of the POS was also performed where needed.

Furthermore, we added another layer of granularity to indels that were misidentified by examining the difference in better positioning the mismatch between the caller output and the ground truth. A decision tree was used for these steps, depicted in **Figure 3**, in order to categorize each indel as diff POS, diff REF and diff ALT if they were mismatched in terms of genomic position, reference allele and alternate allele column of the vcf, respectively.

**Figure 3:**
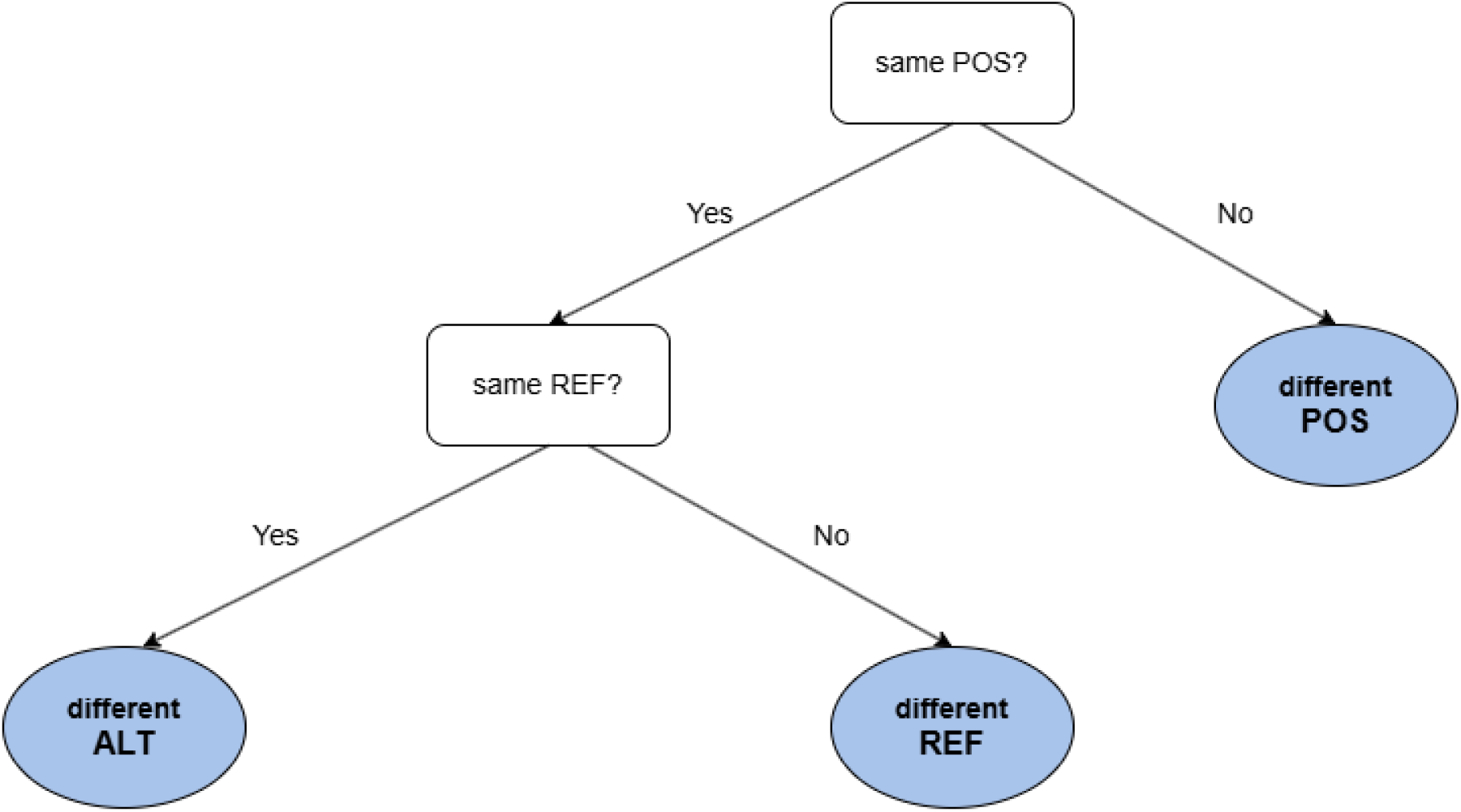
Decision tree for mismatched indels categorization. At each node, variants were compared for identical genomic position (POS), reference allele (REF), and alternate allele (ALT). Variants were classified as differing in POS, REF, or ALT depending on the outcome of these comparisons.

#### 2.2.5 Define classes and metrics

To systematically evaluate the performance of variant callers, we introduced a set of metrics designed to assess their accuracy and reliability in detecting both SNVs and indels. We defined three classes of variants;

- those that were present only in the ground truth but not reported by the variant caller, i.e. the **False Negative** (FN),
- those that were present in the ground truth and also reported by the variant callers, i.e. their overlap or **True Positive** (TP) and finally,
- those that were only reported by the variant caller but did not exist in the ground truth, i.e. **False Positive** (FP).

These classes were applied to all independent analyses of SNVs and indels.

Beyond identifying TP variants, we evaluated the deviation of the callers’ AF estimates from the ground truth. To properly capture this relationship, we defined a new metric that quantifies the deviation between the AF reported by each caller and the AF of the ground truth, as shown in **Equation 3**:

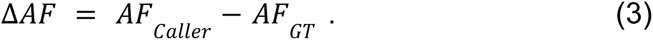

It becomes apparent that the ideal scenario is when the AF deviation equals zero (Δ*AF* = 0), which can be achieved when the reported AF from the callers is the same as the one of the ground truth. On the other hand, Δ*AF* > 0 when a variant caller provides an AF that is higher compared to the real value in the ground truth. Finally, Δ*AF* < 0 represents variants that are falsely reported with lower AF by the callers. It should be noted that this comparison can be only applied in the TP variants; in the case of the FP variants the AF of the ground truth equals zero (*AF*_*GT*_ = 0), while the AF of the caller is zero for FN variants (*AF*_*Caller*_ = 0).

To measure the callers’ performance we implemented the ratio of TP against FN for each individual caller, by using the Recall (aka true positive rate and sensitivity), as shown in **Equation 4**:

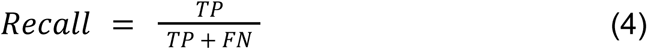

as well as the Precision (aka positive predictive value and specificity), which calculates the fraction of the called variants that were actually TP and is shown in **Equation 5**:

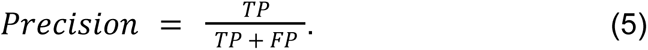

It should be noted that, within the scope of our analysis, no variants fall into the True Negative (TN) category. In our context, TN would correspond to variants absent from both the ground truth and the caller’s output.

## 3. Results: Benchmarking Findings

In this section, we present the findings from our experiments based on the synthetically generated datasets in order to benchmark the variant callers. Results were structured into two sections to allow better insight. First, we provide an overview of callers’ behaviour and performance by implementing grid experiments across the genomic feature space. Second, we focus on variant classes (i.e. TP, FP, FN) from the three independent analyses: SNV true variants, SNV noise and indels. Finally, we report technical aspects related to runtime and memory usage.

### 3.1 Synthetic Datasets

The synthetically generated datasets were designed to explore the effect of different NGS parameters on variant calling. In total, 29 datasets across 620 generated files were studied. More specifically, 25 grid datasets were generated with varying coverage values (300×, 700×, 1000×, 3000× and 5000×), and different read lengths, ranging from 50 bp to 300 bp (50, 75, 100, 150 and 300 bp) to cover all the feature space, depicted in **Table 2**.

**Table 2:** The 25 generated datasets for the benchmarking process. Each row represents a specified value of coverage and each column a specified read length value. All datasets were named based on the corresponding feature values using the convention {coverage}_{read length} (e.g. 1000_150 denotes the dataset with 1000x coverage and 150bp read length).

|  |  | Read Length |  |  |  |  |
| --- | --- | --- | --- | --- | --- | --- |
|  |  | 50bp | 75bp | 100bp | 150bp | 300bp |
| Coverage | 300x | 300_50 | 300_75 | 300_100 | 300_150 | 300_300 |
|  | 700x | 700_50 | 700_75 | 700_100 | 700_150 | 700_300 |
|  | 1000x | 1000_50 | 1000_75 | 1000_100 | 1000_150 | 1000_300 |
|  | 3000x | 3000_50 | 3000_75 | 3000_100 | 3000_150 | 3000_300 |
|  | 5000x | 5000_50 | 5000_75 | 5000_100 | 5000_150 | 5000_300 |

The number of reads ranged on average from ∼100,000 reads, for datasets with lower coverage, to ∼300,000 reads for “deeper” datasets. Finally, four additional datasets were generated to evaluate the effect of the number of individual files on the resulting AF in the merged file, considering several scenarios (i.e., 1, 2, 4, and 10 individual files) prior to implementing the 10-file configuration described in Subsection 2.2.2. All datasets were of high quality (i.e. properly paired reads of high quality with no secondary, duplicates etc.) as they provide the most reliable ground truth for assessing methodological performance.

### 3.2 Callers’ behaviour across the genomic feature space

This section presents an overview of the performance of variant callers across the genomic feature space. To systematically assess their behavior, we performed an ablation study in which individual parameters were subjected to independent variation, while all others were held constant. This approach enabled the evaluation of the influence and significance of each parameter on variant calling.

We studied caller’s **FN variants trends** by studying the recall for all grid experiments. As seen in **Figure 4**, recall values were consistently higher for SNVs than indels and higher for true variants compared to noise. Among callers, VarScan2 achieved the highest recall for true variants, but also showed the largest variability across analyses. VarDict and LoFreq followed, performing similarly in the noise analysis. In contrast, FreeBayes and Mutect2 yielded the lowest recall across true variants, noise and indels.

**Figure 4:**
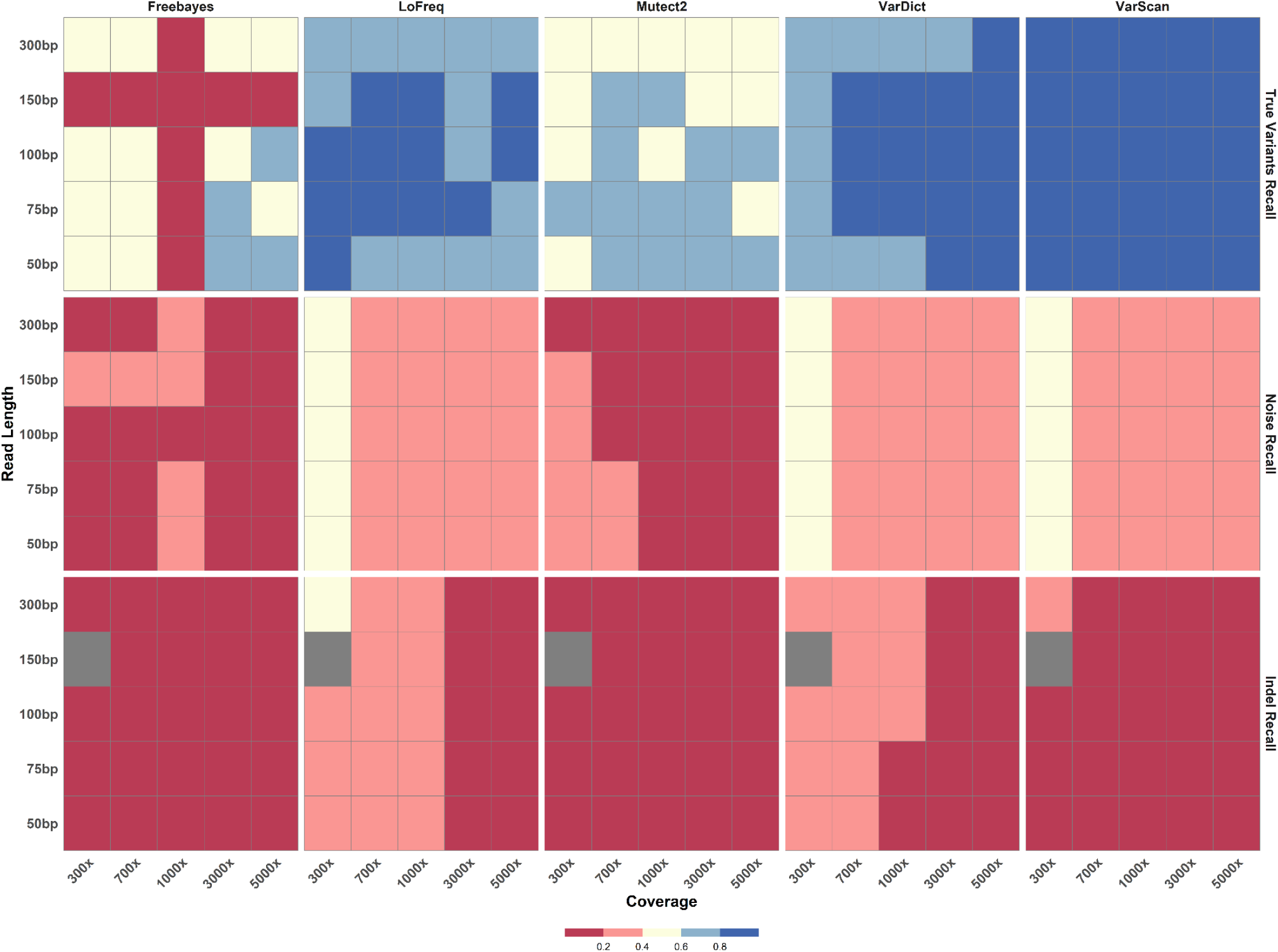
Heatmap showing recall for all experiments across all variant callers. Rows represent read lengths, and columns represent coverage values. Colors indicate recall values, ranging from low (red) to high (blue) in increments of 0.2, with gray indicating missing values. Three independent analyses are presented: top, SNVs True Variants; middle, SNVs Noise; and bottom, Indels. Each panel corresponds to a different variant caller.

Sequencing parameters influenced caller performance in distinct ways. In terms of noise, most callers (LoFreq, Mutect2, VarDict, VarScan2) showed better recall at lower coverages, whereas true variant detection was more dependent on read length. Mutect2 performed best for shorter reads (50–75 bp), while LoFreq and VarDict favored the shortest (50 bp) and longest (300 bp) read lengths. All relevant data for these associations can be found in **Section S4** of the Supplementary Material.

**FP variants trends** were assessed by studying the precision for all grid experiments. As seen in **Figure 5**, precision values were consistently higher for SNVs than indels since most callers did not generate FP variants in the true variant and noise analyses, with the exception of VarScan2 that called a few FP variants at lower coverage values. VarScan2 and Lofreq performed best in the indels analysis, followed by VarDict and Mutect2. All data for this plot can be found in **Section S4** of the Supplementary Material.

**Figure 5:**
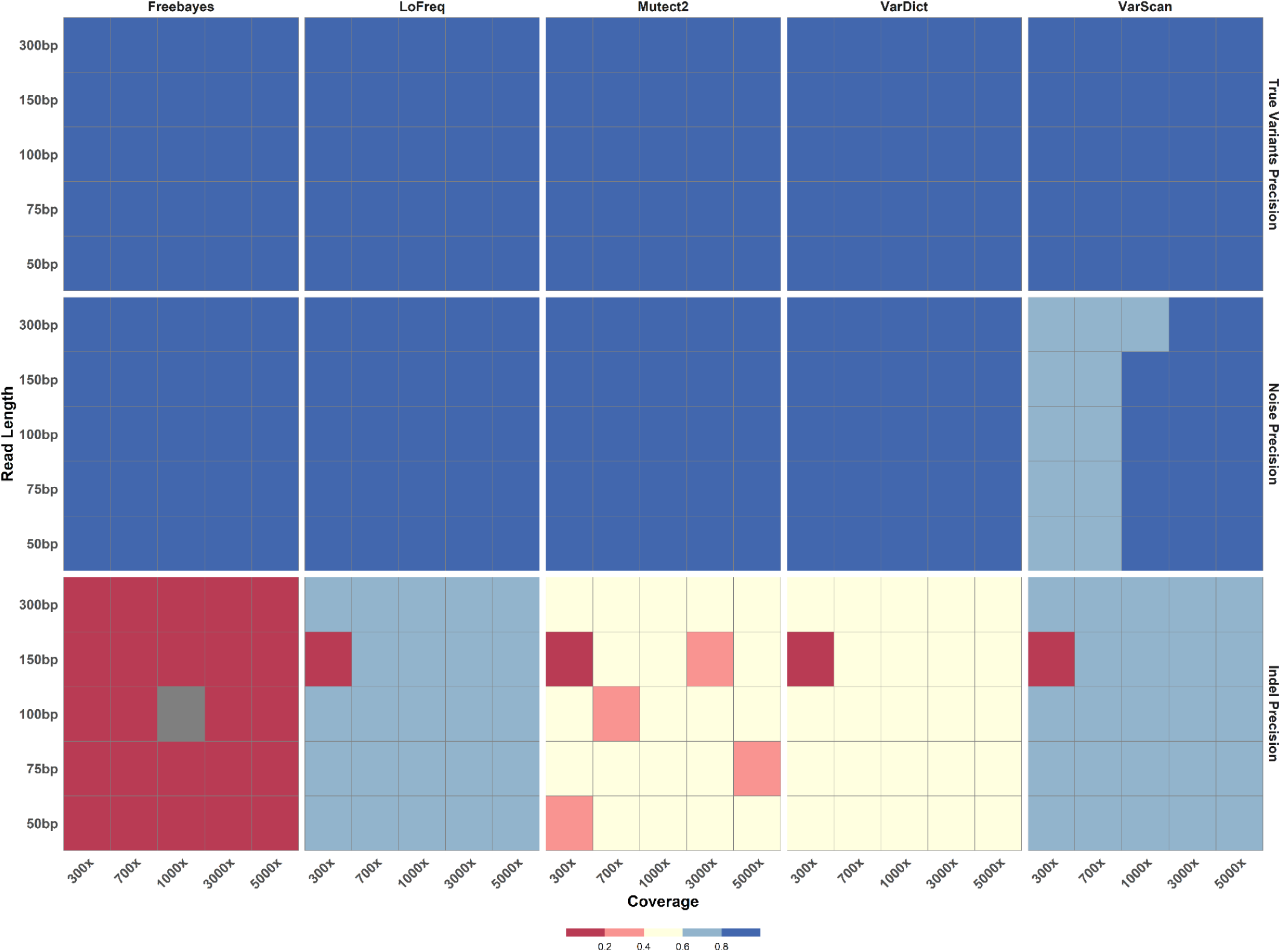
Heatmap showing precision for all experiments across all variant callers. Rows represent varying read lengths, and columns represent coverage values. Colors indicate precision, ranging from low (red) to high (blue) in increments of 0.2, with gray indicating missing values. Three independent analyses are presented: top, SNVs True Variants; middle, SNVs Noise; and bottom, Indels. Each panel corresponds to a different variant caller.

To investigate the frequency resolution of variant callers, we applied an AF binning strategy following a previous study [7], as illustrated in **Figure 6A–I**. Recall and precision metrics are shown for all callers across the different AF bins. True variants were not detected at very low frequencies (AF < 0.02, **Figure 6A-E**),, while indels and noise were present across all frequency ranges. SNVs and indels formed separate clusters in the AF range of 0.005–0.5 (**Figure 6D-H**), with indels displaying a more dispersed distribution within this frequency band. Most variant callers’ SNVs calls clustered tightly near high precision (≈1.0) across all AF bins but exhibited greater variability in recall, particularly at the lowest AF values. As AF increased the points progressively concentrated toward the upper-right corner of the plot, reflecting both high precision and high recall. Notably, indels consistently showed lower recall and broader variability compared to SNVs and noise across all bins. Mutect2 was the only caller that reported variants from all three analyses in the higher frequency band (**Figure 6I**). All relevant data for this analysis can be found in **Section S5** of the Supplementary Material.

**Figure 6:**
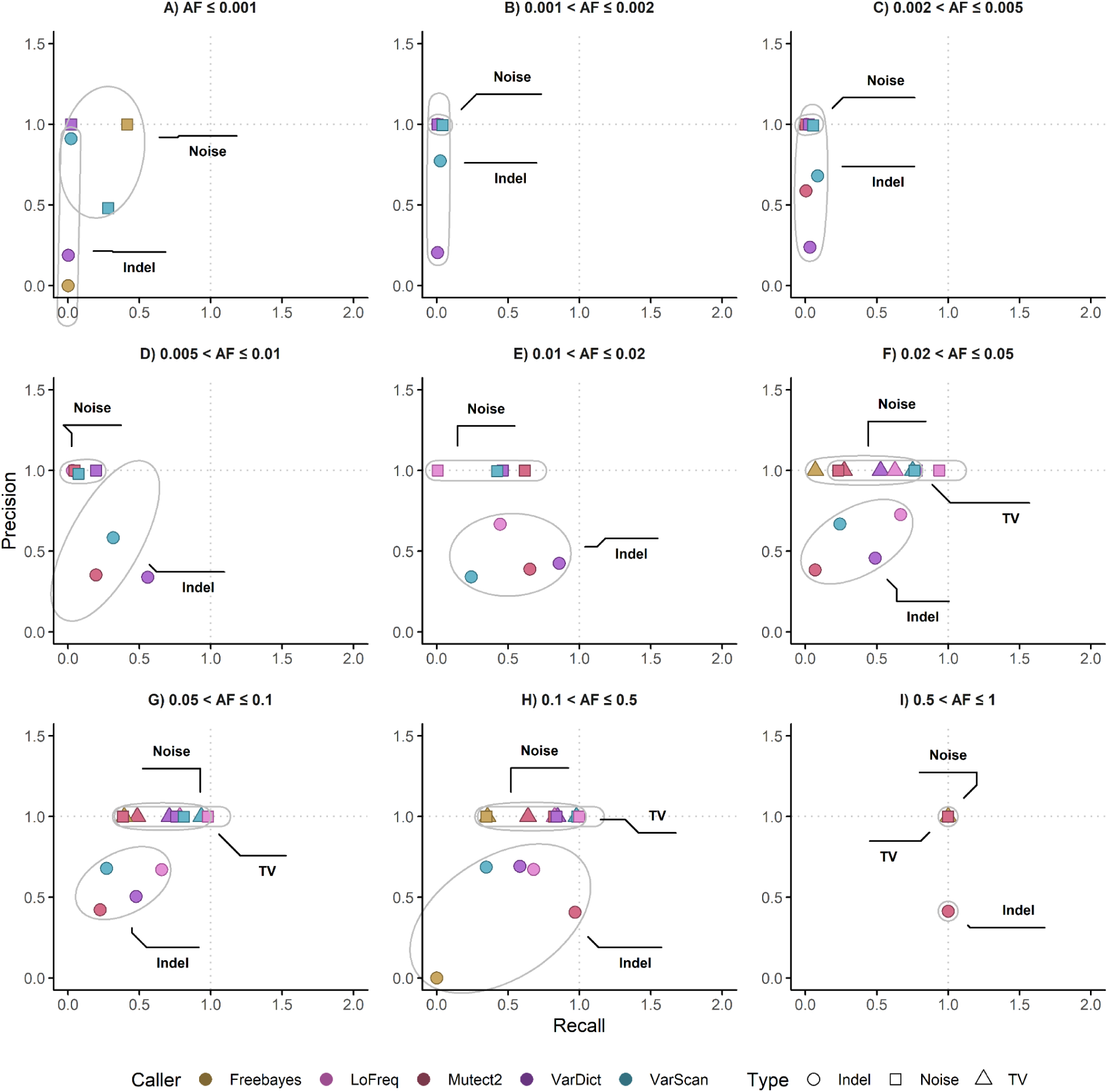
Frequency resolution of variant callers across different AF bins. Recall is plotted on the x-axis and precision on the y-axis. Each color represents a different algorithm, and each marker type corresponds to one of the three independent analyses (Indels, Noise, or True Variants [TV]). Axes are extended slightly beyond the observed metric ranges for visualization, with dashed reference lines at a value of 1 to aid interpretation. Panels correspond to AF ranges: (A) <0.1%, (B) 0.1–0.2%, (C) 0.2–0.5%, (D) 0.5–1%, (E) 1–2%, (F) 2–5%, (G) 5–10%, (H) 10–50%, and (I) 50–100%.

Next, we evaluated how accurately the callers quantified each variant in the SNVs True Variant analysis, focusing specifically on the precision of their reported AF. To this end, we defined a new metric, the AF deviation (ΔAF), which quantifies the difference between the reported AF and the ground truth. As shown in **Figure 7A**, LoFreq and VarDict consistently demonstrated the most stable and accurate performance, with a near-zero mean error across all combinations of read length and coverage. The uniform dark blue color of their respective heatmaps indicates a high degree of accuracy and robustness to changes in sequencing parameters. In stark contrast, Mutect2 showed the greatest deviation in its mean AF estimation, showing a significant positive bias (overestimation) at longer read lengths, particularly for lower coverage values. It is also worth noting that Mutect2 had the highest ΔAF dispersion, by reaching both the highest values and the lowest values (0.87 and -0.26, respectively). On the contrary, LoFreq exhibited the best performance with its highest value being 5. 1 × 10^−7^ and the lower -0.019, as shown in **Section S7** of the Supplementary Material. FreeBayes and VarScan2 performed generally well, but with slight biases (FreeBayes for shortest reads and VarScan2 for longer reads), suggesting less consistent accuracy compared to VarDict and LoFreq.

**Figure 7:**
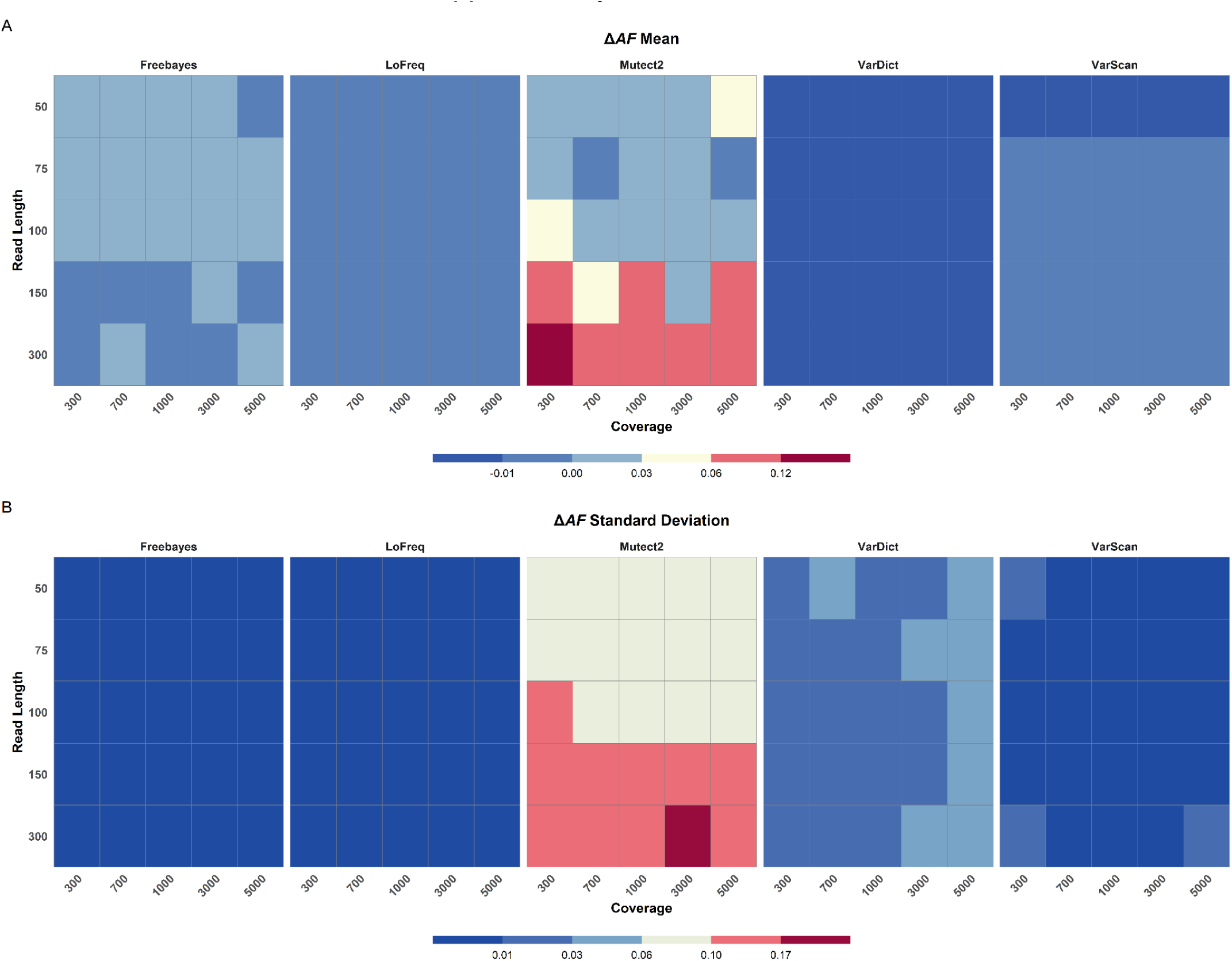
Heatmap showing ΔAF, as defined in Equation 3, for the SNVs True Variants analysis. Rows represent varying read lengths, and columns represent different coverage values. Colors indicate mean and standard deviation values, ranging from low (best, blue) to high (worst, red); note that the scale and range of each legend differ for better visualization. **(A)** Mean deviation values are shown, and **(B)** standard deviation values are depicted.

The ΔAF standard deviation is depicted in **Figure 7B**. FreeBayes, LoFreq and VarScan2 showed consistently low standard deviations across the entire feature space. Conversely, Mutect2 displayed the highest standard deviation, particularly for higher read lengths and coverages. VarDict also showed some variation, but its standard deviation remained lower than that of Mutect2 across all conditions. All data for these analyses can be found in **Section S6** of the Supplementary Material.

### 3.3 Characteristics of variant classes (TP, FP, FN)

This section focuses on the different classes of variants, with the aim of disentangling how caller performance is shaped by variant type. To this end, the results from each independent analysis were systematically aggregated and compared, allowing us to identify characteristic trends, recurrent patterns and distinctive features associated with each variant class. This comparative perspective provides deeper insights into the underlying behavior of the callers, highlighting both commonalities and divergences in their capacity to detect and accurately characterize different classes of variants.

We examined variant classes by focusing on SNV true variants, aggregating results across all datasets and callers. **Figure 8A** shows broadly similar AF distributional shapes for FNs and TPs, while **Figure 8B** summarizes counts by caller. VarScan2 yielded the most TPs, followed by VarDict and LoFreq. All callers reported some FNs, however, VarScan2 recorded the fewest (272), with LoFreq being the next best choice with 1,311 FNs. Both Mutect2 and FreeBayes showed higher numbers of FN variants. As seen in **Figure 8C**, which presents ΔAF densities, Mutect2 exhibited the largest dispersion around the mean. VarScan2’s density peaked at negative ΔAF and VarDict showed a primary negative peak with a secondary peak at lower ΔAF. By contrast, LoFreq and FreeBayes outputs remained tightly centered near zero.

**Figure 8:**
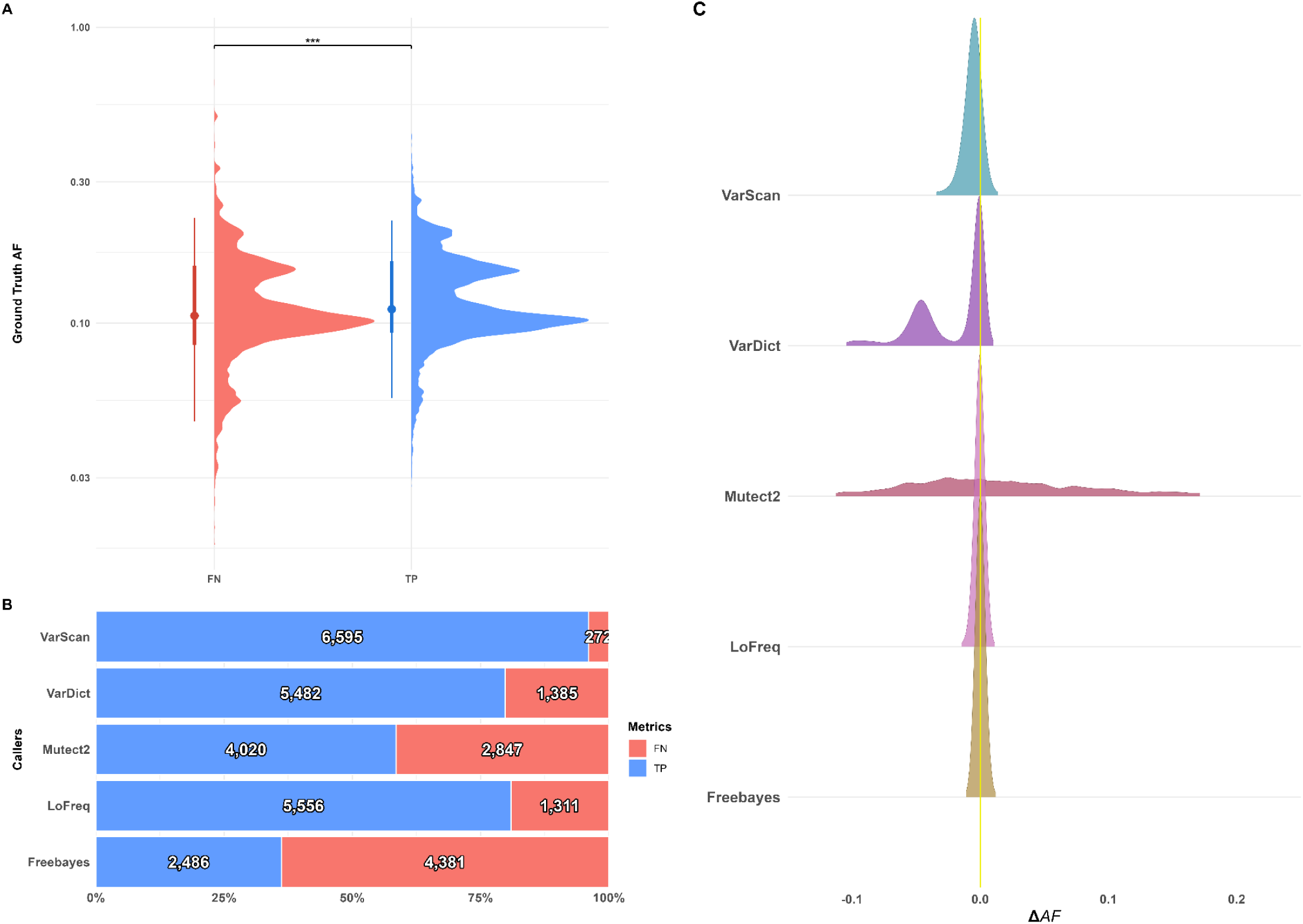
Multiplot presenting a comprehensive benchmark of variant classes (TP, FN, FP) for the SNVs True Variants analysis. (A) Violin plots showing the distribution of ground truth AF for TP variants (blue) and FN variants (red). (B) Stacked bar charts quantifying the number of TP (blue) and FN (red) variants for all callers; bars are normalized to 100% to facilitate direct comparison of TP and FN proportions, with absolute counts displayed on each bar. (C) Density plots of ΔAF.

As seen in **Figure 9A**, LoFreq, VarDict and VarScan2 closely “mimicked” the Ground Truth DP distribution. Mutect2 was the outlier, with its median DP being well below the Ground Truth. A slight shift towards lower coverage values was also seen for VarScan2. **Figure 9B** shows that LoFreq and FreeBayes were well aligned with the Ground Truth, while VarScan2 and VarDict showed a systematic DP underestimation. Finally, Mutect2 exhibited the widest AF dispersion with its median values being above the mean of the ground truth.

**Figure 9:**
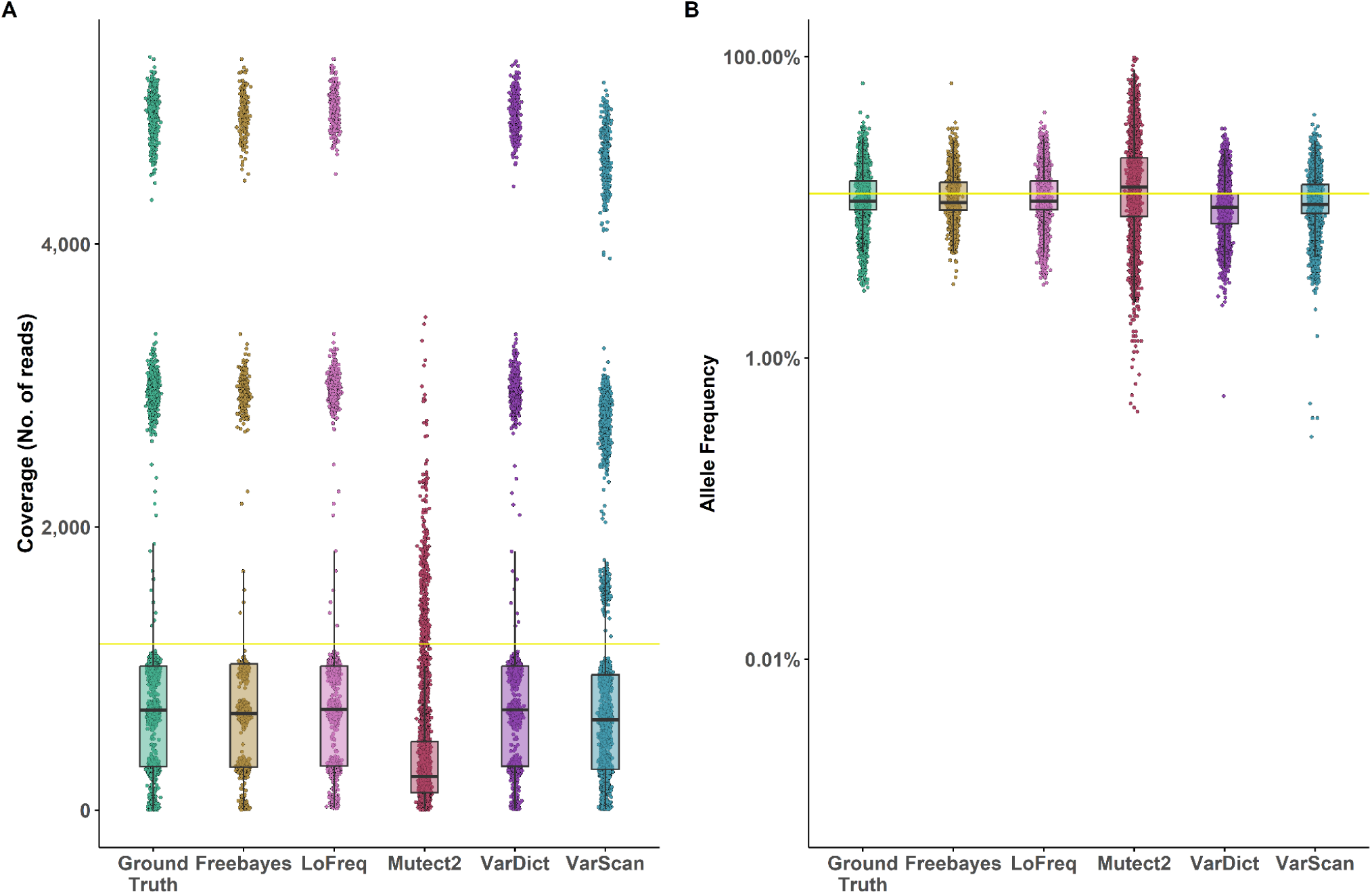
Multipanel figure illustrating the performance of all variant callers for SNVs True Variants with respect to coverage and AF compared to the ground truth. (A) Boxplots of coverage for all true variants in the ground truth and for each caller; each dot represents an individual variant. The yellow line indicates the mean coverage of the ground truth. (B) Corresponding AF boxplots for the same TP variants for each caller. The y-axis is logarithmic to better display the distribution of AF values across orders of magnitude. The yellow line indicates the mean AF in the ground truth. Each caller is shown in a distinct color, consistent across panels to facilitate direct comparison.

We analyzed variant classes by focusing on SNV noise, aggregating results across all datasets and callers. As seen in **Figure 10A**, AF distributions were clearly distinguished by class with FNs being concentrated at the lowest band, FPs occupying an intermediate AF band and TPs showing the highest values. **Figure 10B** depicts that this pattern actually mirrors AF: FNs and FPs had very low alt depths (AD) with TPs being characterized by substantially higher AD. Class distributions were dominated by FNs across callers, as can be seen in **Figure 10C**. At the level of individual callers, VarScan2 reported the most TPs (∼378k) but also carried a non-trivial FP burden (∼81k). LoFreq and VarDict reached intermediate TP efficiency (∼340k and ∼282k respectively), while Mutect2 reported few FPs (74) and a lower TP count (∼161k). Finally, FreeBayes recorded the fewest TPs (∼158k) and the highest number of FN variants (∼909k).

**Figure 10:**
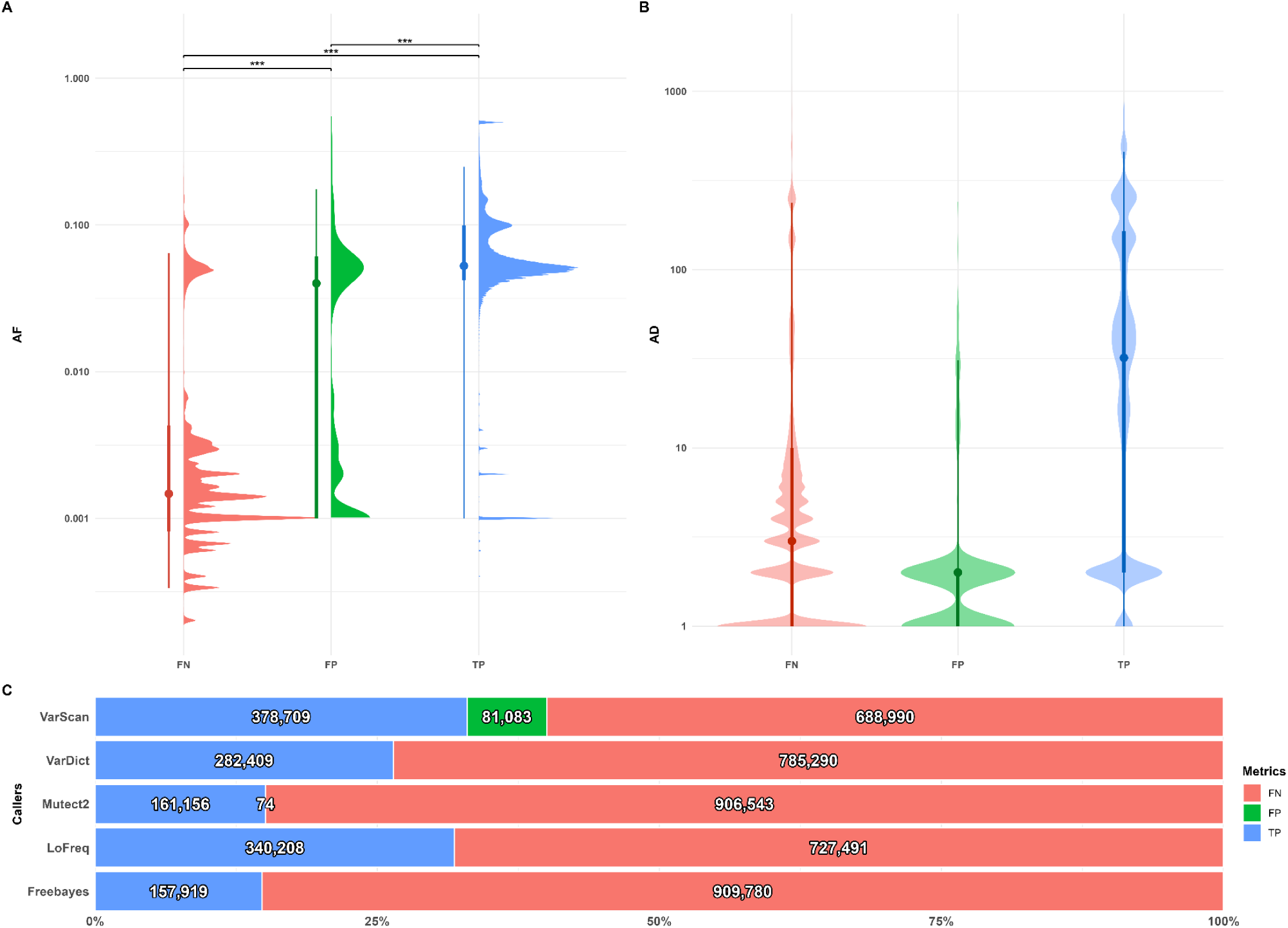
Multiplot presenting a comprehensive benchmark of variant classes (TP, FN, FP) for the SNVs Noise analysis. (A) Violin plots of AF for FN (red), FP (green), and TP (blue) variants across all callers; the y-axis is logarithmic. (B) Violin plots of alternate allele depth (AD) for FN, FP, and TP variants on a logarithmic scale. (C) Stacked bar charts quantifying TP (blue) and FN (red) variants for all callers; bars are normalized to 100% to facilitate direct comparison of TP and FN proportions, with absolute counts displayed on each bar.

Finally, we integrated all results and data from the indel analysis. As shown in **Figure 11A–B**, indel FNs were concentrated at the lowest AF and AD, while FPs and TPs had the highest values with more narrow spreads. Regarding caller performance, depicted in **Figure 11C–D**, indel detection showed low recall across all callers, with varying precision; LoFreq and VarScan2 reached a Precision≈0.70 but recall remained limited (≈0.20 and ≈0.09, respectively). VarDict attained Precision≈0.46 and recall≈0.15, while Mutect2 showed Precision ≈0.41 and recall ≈0.06. FreeBayes reported FPs without TPs (Precision = 0, recall = 0). As seen in **Figure 11D**, FNs dominated every caller report, with LoFreq and VarDict yielding the largest TP totals, VarScan2 recovering fewer TPs than those two, and Mutect2 and FreeBayes contributing the fewest TPs. **Figure11E** offers insights regarding the indels subcategories, as defined in **subsetion 2.2.4**, showing that most mismatches arise from positional discordance (diff POS) followed by diff ALT category, across all callers. As depicted in **Figure 11F**, the main discordance for FN indels is diff POS, followed by diff ALT category. On the other hand, the most populated category for FP variants is that of diff REF followed by the diff POS category.

**Figure 11:**
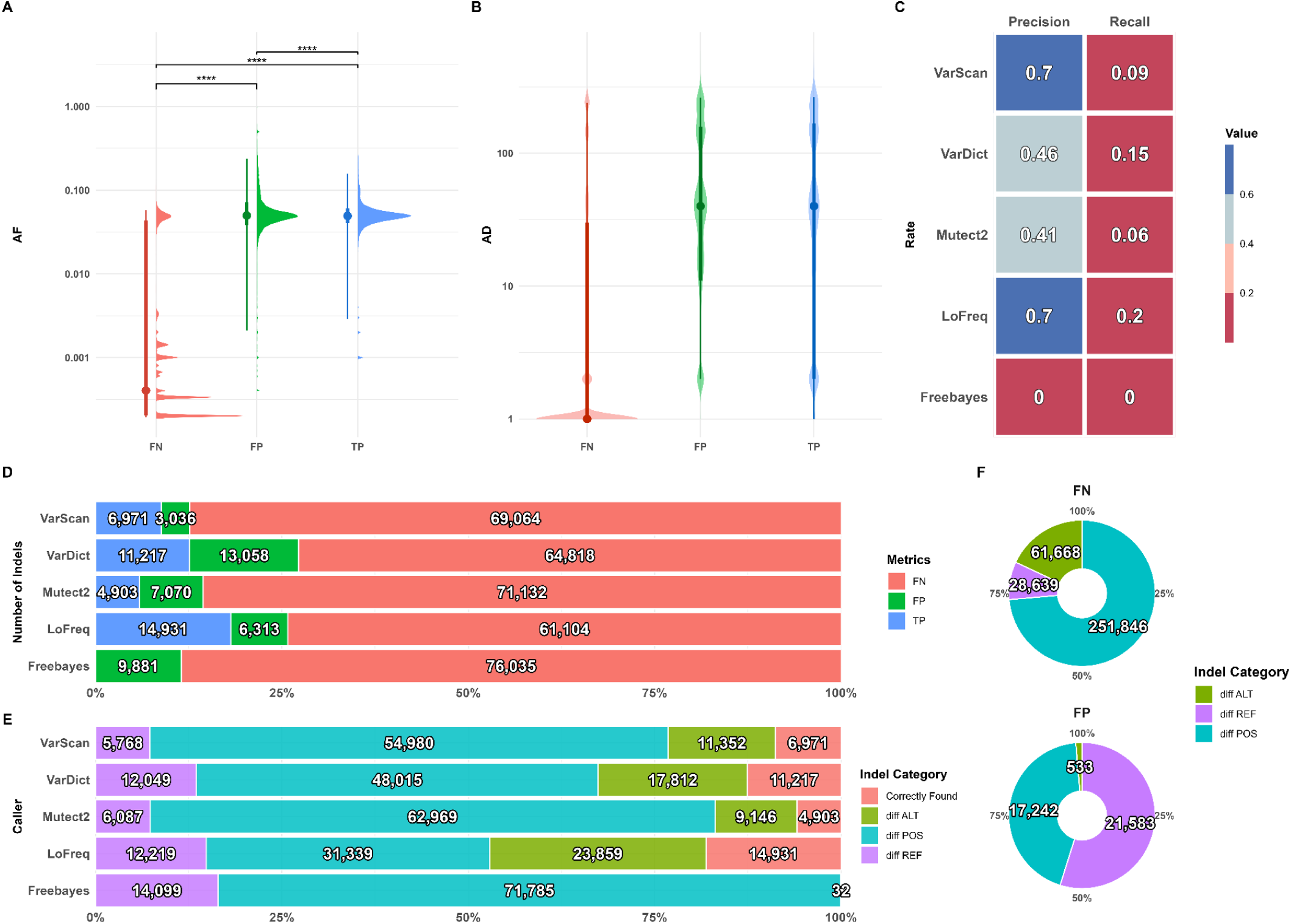
Multiplot presenting a comprehensive benchmark of variant classes (TP, FN, FP) for the indel analysis. (A) Violin plots of allele frequency (AF) for FN (red), FP (green), and TP (blue) variants across all callers; the y-axis is logarithmic. (B) Violin plots of alternate allele depth (AD) for FN, FP, and TP variants on a logarithmic scale. (C) Precision and recall heatmaps for all callers. (D) Stacked bar charts quantifying TP (blue) and FN (red) variants for each caller; bars are normalized to 100% to allow direct comparison of proportions, with absolute counts displayed on each bar. (E) Stacked bar charts showing the number of indels in the categories defined in Subsection 2.2.3. (F) Donut charts summarizing FN (top) and FP (bottom) variant composition across the diff POS, diff REF, and diff ALT categories.

### 3.4 Runtime and Computational Efficiency

A runtime evaluation of synth4bench was also performed. We calculated the total runtime for the generation and variant calling process for one of the most computationally heavy datasets (i.e. that of 5000x coverage, which contained a total of 620,642 reads). The dataset generation step required approximately 6 minutes. For the variant calling stage, the elapsed times were as follows: Mutect2 - 50 minutes, VarDict - 12 minutes, LoFreq - 12 minutes VarScan2 - 3 minutes and FreeBayes - 2 minutes.

Furthermore, we recorded the time needed for our R scripts to execute the benchmarking as well as the visualization for 10 datasets. As expected, processing time increased with the number of reads generated at higher coverage levels with a mean *S4BR.R* run time of 25.22sec. On average, the visualization script (*S4BR_plot.R*) required more time to be executed (equal to 39.1 sec) and was not affected by dataset depth. Values for the run times comparison are presented in **Section S3** of the Supplementary Material.

All experiments were performed on a desktop system equipped with an Intel(R) Xeon(R) CPU E5-2673 v3 @ 2.40GHz and 32 GB of RAM, running in WSL2 within a conda environment.

## 4. Discussion: Lessons Learned

Here, we present a novel pipeline, synth4bench, for benchmarking somatic variant callers through the use of synthetic genomics data as ground truth. Our approach showed that callers do not behave similarly due to their algorithmic nature that weighs the involved NGS parameters differently. Based on this, we developed synth4bench to shed light on the underlying principles that govern the performance of each variant caller by analyzing distinct datasets across the NGS feature space and observing each caller’s response to the range of the applied parameters.

We mainly focused on the **technical evaluation** of variant callers, hence variant annotation was not incorporated and all chromosomal positions were treated equally. While biological context, such as certain genomic regions or hotspots, is undoubtedly important, our approach deliberately isolated methodological performance, allowing the observed differences between callers to reflect technical factors.

Our analysis revealed distinct patterns in **recall and precision** performance across variant callers, variant types and NGS configurations. The higher recall for SNVs compared to indels and for true variants compared to noise, is consistent with known challenges in indel detection and highlights the greater reliability of callers for SNVs [8,9,15,16,45–47]. The dependence of recall on NGS parameters further emphasizes the interplay between algorithmic design and data characteristics.

It was observed that read length had a more decisive effect on true variant detection (**Figure 4**). The absence of FP for true variants and noise (**Figure 5**) can be largely attributed to the way these analyses were constructed. By defining all non-true variants as noise, the framework effectively eliminated scenarios where FP could occur. This design highlights the limitations of interpreting absolute precision values in isolation, as they may reflect dataset composition rather than caller performance alone.

The joint assessment of recall and precision across AF bins (**Figure 6**) provides important insights into callers’ **AF resolution**. Within the intermediate range (between 0.005-0.5) (**Figure 6D-H**) SNVs and indels became more distinguishable; SNVs showed tighter clustering, while indels remained more dispersed, consistent with their known detection challenges [8,9,15,16,45–47]. This separation of indels from SNVs in the frequency spectrum indicates that indel calling remains more error-prone and sensitive to AF, reflecting inherent algorithmic limitations in handling indels at low frequencies. As AF increased, recall improved notably for all three analyses, reflecting the combination of high precision and high recall. Importantly, indels consistently underperformed SNVs in recall, confirming that they remain the most challenging variant type across AF ranges [8,9,15,16,45–47]. Improvements in variant detection with increasing AF confirm that caller performance becomes more stable as variant support strengthens, something also visible in noise analysis results showing that FNs and FPs clustered towards the lower AF spectrum, while TPs had substantially higher AF (**Figure 10B**). Based on our findings, callers like VarScan2 and VarDict may be more suitable for variant detection at the lower end of the AF spectrum.

When analysing indels, even when precision is moderate to high for some callers, recall is uniformly low, driven by FNs at very low AF; As indicated in **Figure 11C**, almost all callers achieved better precision values than recall. Our analysis offers extra granularity regarding indel mismatches, revealing distinct patterns for detection failures versus false reports. FN indels (i.e. missed variants) were predominantly caused by errors in chromosomal positional placement (diff POS) relative to the ground truth. Conversely, FP indels (i.e. ‘imagined’ variants) were characterized by errors stemming from both different reference allele (diff REF) and chromosomal positional inaccuracies (diff POS) (**Figure 11F**). The choice of caller should therefore reflect the study aim, with precision prioritizing workflows leading to better results when implementing LoFreq and VarScan2, whereas recall-critical prioritizing workflows being more efficient when integrating VarDict, which had the highest recall values and the smallest precision and recall trade-off. Some use cases may require caller ensembles, realignment-aware post-processing and lower evidence thresholds coupled with artifact filters to recover more true indels without inflating FPs.

Regarding true variants, our results indicate that adequate candidates for their detection are VarScan2 for maximizing TP recovery, but with awareness of its mild AF underestimation and its tendency to produce FP variants, and LoFreq when precise AF estimation is paramount.

The performance of **Freebayes** in our somatic variant benchmark was notably constrained, yielding the poorest overall results among the five callers tested. This outcome is consistent with the fact that Freebayes was not initially developed for somatic variant calling, instead employing a Bayesian, haplotype-based framework that models genotype likelihoods and infers ploidy across a cohort [34,49]. The limitations of this methodology were evident across several metrics; the primary source of discordance was systematic misrepresentation of the REF allele in its VCF output, especially for indels. Furthermore, Freebayes struggled with variant type complexity and low-frequency variants. In particular, the algorithm failed to recover any TP indels, contributing only to FP and FN for indels, while significantly underperforming at the detection of variants with low AF. This consistent low recall underscores its limitations in sensitivity under the tested conditions (**Figure 4**), that predominantly included somatic mutations with low AF. Despite issues with accuracy and sensitivity, Freebayes demonstrated stable AF estimation precision. Its standard deviation of the ΔAF was consistently low (**Figure 8C**), indicating high precision. However, during the study of ΔAF across varying read lengths and coverage (**Figure 7**), Freebayes showed slight biases and variations under certain conditions. While not as dramatic as those observed for other callers, these variations suggest that Freebayes is less robust to changes in sequencing parameters than LoFreq or VarDict. The stability in AF estimation precision, coupled with a high number of FN variants, is a clear indication that Freebayes tends to be conservative, preferring to make fewer, more precise calls, which result in reduced recall (**Figure 4**).

The performance of **Mutect2** presented a complex picture in our benchmark, exhibiting behaviors that both align with and contrast with previous studies. Some reports highlight its capacity to detect SNVs at low coverage and AF [14], however, our observations indicated a consistent low recall (**Figure 4**), underscoring a limitation in sensitivity. Mutect2 frames somatic detection as a Bayesian hypothesis test on allele fraction, running two models (reference *M*_0_ and variant 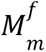) to distinguish true variants from sequencing errors [33]. Similar to LoFreq, it assumes independent base errors at different loci. A key challenge noted in prior work is Mutect2’s tendency to produce a substantial number of FP calls in its tumor-only mode [8], a common issue for methods that rely on filtering germline mutations using population databases [50]. Our findings indicate distinct issues concerning data characteristics. Mutect2 performed better for shorter read lengths, but showed significant vulnerability to longer reads and higher coverages (**Figure 7**). This behavior manifested as a pronounced bias and lack of precision in AF estimation under these specific conditions, suggesting its performance is highly sensitive to the exact combination of NGS parameters, a critical design consideration. For indels, Mutect2 was notably conservative, exhibiting low precision across the entire AF spectrum. While its recall increased as AF reached higher values, its overall sensitivity remained constrained. Furthermore, Mutect2’s reported AF estimates were consistently the noisiest, as reflected by the greatest AF variability (**Figure 8C**). This variance is primarily attributed to Mutect2’s internal downsampling of reads (**Figure 9A–B**). This process reduces the effective depth used for variant calling, regardless of the dataset original depth, leading to two major consequences: a markedly lower reported DP compared to Ground Truth and a widening of the AF dispersion. This reduction in effective depth increases the variance of AF estimates, which can negatively impact recall, especially for variants near the detection limit even in datasets with high available coverage.

**VarScan2** operates fundamentally differently from the haplotype-based callers, relying on a pileup file as input that allows to “see everything” at all genomic positions before applying a series of ad hoc filters and thresholds to generate the final variant list [36]. When a paired normal sample is provided, it implements a Fisher exact test to statistically compare the two input files and discern somatic variants. This approach yielded mixed, yet generally robust results in our benchmark. VarScan2 demonstrated remarkable capabilities in detecting both indels and low-frequency SNVs, being one of the few callers able to report variants with an AF below 0.1% (**Figure 6**). Its strong overall recall (**Figure 4**) suggests it can be a highly effective tool for true variant detection. For indels, it exhibited higher precision than some other callers but still missed a significant number of true indels, indicating a limitation in its sensitivity for this variant type. Regarding AF estimation, VarScan2’s performance was generally good (**Figure 7 and 8C**). However, its ΔAF distributions often showed a slight negative-centered bias (**Figure 8C**), indicating a tendency to underestimate AF. This suggests that while it is robust, it may be less stable or accurate than algorithms like LoFreq or VarDict under varying read length and coverage conditions. A particularly noteworthy finding came from the noise analysis (**Figure 10B**), where VarScan2 successfully captured the most TP noise variants compared to all other algorithms. This further underscores its sensitivity, making it suitable for identifying true variants.

The performance of **LoFreq** in our benchmark demonstrated exceptional robustness and precision, aligning with its established strengths in detecting rare variants [35] and its capacity to discern TP reported in previous studies [7]. LoFreq’s core methodology is rooted in a null hypothesis that posits all variants arising from sequencing error. It models the presence of a variant as a series of independent Bernoulli trials, with each base’s error probability directly informed by its Phred quality score. This heavy reliance on quality scores makes our well-characterized benchmark an ideal scenario for studying LoFreq. This methodology, however, represents both a strength and a potential weakness, as it introduces an inherent bias [35] due to the ideal, high quality reads used in our synthetic dataset. Despite this, LoFreq’s practical performance was highly favorable. Its ability to call rare variants was confirmed, and its consistent accuracy and precision in AF estimation across the entire feature space (**Figure 7**) demonstrated remarkable robustness against variations in read length and sequencing coverage. Concerning indels, LoFreq was the best-performing caller, yielding the largest TP totals, while also achieving the highest precision (**Figure 11**). Although it still missed some true indels, its overall performance in this challenging category was superior. For overall recall (**Figure 4**) LoFreq was moderately affected by the read length, with a slight recall decrease at extreme lengths (50 bp and 300 bp). Finally, in the noise analysis (**Figure 10B**), LoFreq was the second most successful algorithm, after VarScan2, at capturing TP noise variants. While we did not observe the large numbers of FP reported in some prior studies [4,7,8], likely a result of our data generation methodology taking as noise all low-AF variants that were excluded from the true variant analysis as explained in **Section 2.2.3**, LoFreq’s high precision suggests it effectively manages to detect low-frequency variants. This overall profile establishes LoFreq as one of the most reliable and technology-independent callers for robust AF estimation across diverse sequencing conditions.

**VarDict** implements a robust strategy for variant calling that significantly improves sensitivity by performing local re-alignment. As detailed in previous work [37], VarDict leverages read information by treating mismatches and soft-clips as potential indel clues. By locally re-aligning read ends (a supervised approach) and mining shared soft-clip sites that flank potential indel regions (unsupervised), it effectively recovers hidden evidence and reports more accurate AFs. This methodological approach contributed to VarDict’s strong performance in AF estimation. During the study of ΔAF (**Figure 7 and 8C**), VarDict demonstrated exceptional robustness and reliability, exhibiting highly consistent accuracy and precision across all genomic feature space. However, a slight tendency to underestimate AF was noted, with its ΔAF distributions showing a second peak at negative values indicating a bias (**Figure 8C**). For overall recall (**Figure 4**) similarly to LoFreq, its performance was also moderately affected by the read length, with a slight recall decrease at extreme lengths (50 bp and 300 bp). VarDict’s local re-alignment strategy was particularly impactful for indels. It ranked second for TP indel recovery and achieved the highest overall indel recall (albeit a modest ∼0.15) by effectively minimizing FN.

Our findings align with what the extensive literature documents, which is the **persistent disagreement** among different variant callers, with studies frequently showing little to no overlap in results even when analyzing the same samples [14,16,51]. Beyond this inter-caller variability, outcomes across different benchmarking studies also exhibit substantial inconsistency. This highlights a strong dependence of results on the specific input data used, a challenge that persists even within advanced ML-based ensemble approaches [16,52]. Such pronounced inconsistencies suggest a fundamental limitation, that current algorithms do not yet capture mutational mechanisms and NGS artifacts with adequate fidelity, indicating that fully modeling of the underlying biological and technical processes remains an open challenge.

## 5. Conclusion

In summary, this work provides a systematic technical benchmark of widely used variant calling algorithms, focusing on their ability to detect both SNVs and indels under controlled, high-quality conditions as offered by the generation of synthetic data. Our findings highlight substantial **discrepancies** among callers and even benchmarks, consistent with previous reports and reinforce the notion that existing methods do not yet fully capture the complexity of mutational mechanisms, leaving this an important open challenge for the field. **Indels** remain the hardest variant type to call, and tool selection must be tailored, favoring LoFreq or VarScan2 for precision-focused indel calling or VarDict when recall is prioritized. The trade-off between recall and precision is most acute at low AF, with VarScan2 and VarDict demonstrating suitability for variant detection at this challenging lower end of the AF **spectrum**. For **true SNV variants**, VarScan2 is optimal for maximizing TP recovery, while LoFreq is recommended when precise AF estimation is paramount. Ultimately, achieving maximal **sensitivity and reliability** necessitates caller-specific optimization of sequencing strategies, with an informed choice of algorithm based directly on the study’s priorities.

Despite all efforts, this systematic benchmark has potential pitfalls. In relation to the **quality** of the data used for evaluation, we deliberately restricted our analyses to high quality datasets, as they provide the most reliable ground truth for assessing methodological performance, since inherent noise and inconsistencies from noisy data could confound the evaluation and obscure genuine differences between variant calling methods. Nonetheless, extending future benchmarks to include lower-quality data would be valuable, as it could illuminate how variant callers handle actual NGS errors and uncertainties, thereby informing strategies to improve their robustness in real-world applications. Furthermore, there is the aspect of the quality of synthetic data itself. There are various evaluation metrics with respect to fidelity, privacy and applicability, which have been well-addressed by an organized effort by the ELIXIR AI Ecosystem Focus Group [53]. Furthermore, this work focuses primarily on variant calling itself not including read **alignment**, since a previous study [9] demonstrated that the choice of variant caller exerts a greater influence on the detection of SNVs and indels than the choice of alignment algorithm.

As already argued by [19] due to the astronomical number of potential parameter combinations, our study necessarily focused on a fixed set of parameters for each caller according to their best practices, although none proposed specific tumor-only mode instructions. Fully exploring the entire parameter space for every algorithm, an endeavor requiring countless ablation studies, was beyond the scope of this work and also computationally prohibitive.

Finally, from a more **high-level perspective** on the use of synthetic data, it is important to recognize that, despite their popularity in the life sciences, certain limitations must be carefully considered. Synthetic data are highly valuable for the development of new tools as well as for benchmarking existing ones. They are relatively easy to generate or obtain and can be applied to extreme or unusual scenarios that would be prohibitively expensive or even impossible to reproduce in a wet-lab setting. Nevertheless, synthetic data should be regarded as a means to an end, since the ultimate objective is to evaluate hypotheses against real data, which remain the only reliable benchmark of a sound methodological approach.

## Conflict of interest

No conflicts of interest to declare.

## Funding

This work has received funding from **SYNTHIA** - Synthetic Data Generation framework for integrated validation of use cases and AI healthcare applications - is funded by the Innovative Health Initiative Joint Undertaking (IHI JU) under grant agreement No 101172872. The JU receives support from the European Union’s Horizon Europe research and innovation programme and COCIR, EFPIA, EuropaBio, MedTech Europe, and Vaccines Europe and DNV. Funded by the European Union, the private members, and those contributing partners of the IHI JU. Views and opinions expressed are however those of the author(s) only and do not necessarily reflect those of the aforementioned parties. Neither of the aforementioned parties can be held responsible for them This work was also supported by **ELIXIR**, the research infrastructure for life-science data.

## Key actions

- **Indels:** the choice of tool should therefore reflect the study aim, precision prioritizing workflows can prefer LoFreq and VarScan2, whereas recall-critical prioritizing workflows can prefer VarDict which had the highest recall values and the smallest precision and recall trade-off. Some use cases may require caller ensembles, realignment-aware post-processing and lower evidence thresholds coupled with artifact filters to recover more true indels without inflating FPs.
- **Recall and precision:** These findings indicate that sequencing-specific optimization of the selected caller is necessary to maximize sensitivity and reliability, particularly in applications where recall or precision are critical. Caller should be fit-for-purpose, as there isn’t a one-solution-fit-all.
- **Frequency resolution**: The results highlight the trade-off between recall and precision, particularly at low AF ranges where variant detection is most challenging. Callers like VarScan2 and VarDict are more suitable for variant detection at the lower end of the spectrum.
- **True variants:** callers should be selected according to study priorities: VarScan2 for maximizing TP recovery of true variants, with awareness of its mild AF underestimation, LoFreq when precise AF estimation is paramount.
- **Disagreement:** Such pronounced inconsistencies suggest that current algorithms do not yet capture mutational mechanisms adequately, indicating that fully modeling of the underlying processes remains an open challenge.

## Notes

https://zenodo.org/records/16524193

### Competing Interest Statement

The authors have declared no competing interest.

### Summary of Updates

The synthetic data generation pipeline was significantly updated to include Indels (insertions and deletions), which are now thoroughly explored in the variant caller benchmarking. More synthetic datasets were generated and explored under a systematic process, covering a wider range of sequencing aspects. Availability links were updated for the source code repository and the new Zenodo record containing the synthetic datasets. The Methods section was significantly expanded to fully detail the benchmarking process. The downstream analysis was refined and explicitly defined to consist of three independent analyses: two for SNVs (True Variants and Noise) and one dedicated to Indels. A new Metrics section (2.2.5) was introduced, clearly defining performance metrics for assessing accuracy. The Discussion section was substantially revised to incorporate our new findings, offering a more thorough interpretation. The Title and Abstract were revised for clarity and to incorporate specific analytical results. Also, a Graphical Abstract was added to the second page of the manuscript.

## References

1. Zhang L, Jia Z, Mao F et al. Whole-exome sequencing identifies a somatic missense mutation of NBN in clear cell sarcoma of the salivary gland. Oncol Rep 2016;35:3349–56.

2. Martínez-Jiménez F, Muiños F, Sentís I et al. A compendium of mutational cancer driver genes. Nat Rev Cancer 2020;20:555–72.

3. Shin H-T, Choi Y-L, Yun JW et al. Prevalence and detection of low-allele-fraction variants in clinical cancer samples. Nat Commun 2017;8:1377.

4. Olson ND, Wagner J, Dwarshuis N et al. Variant calling and benchmarking in an era of complete human genome sequences. Nat Rev Genet 2023, DOI: 10.1038/s41576-023-00590-0.

5. Li H, Meng L, Wang H et al. Precise identification of somatic and germline variants in the absence of matched normal samples. Brief Bioinform 2024;26:bbae677.

6. Guille A, Adélaïde J, Finetti P et al. A benchmarking study of individual somatic variant callers and voting-based ensembles for whole-exome sequencing. Brief Bioinform 2024;26:bbae697.

7. Xiang X, Lu B, Song D et al. Evaluating the performance of low-frequency variant calling tools for the detection of variants from short-read deep sequencing data. Sci Rep 2023;13:20444.

8. Ha Y-J, Kang S, Kim J et al. Comprehensive benchmarking and guidelines of mosaic variant calling strategies. Nat Methods 2023;20:2058–67.

9. Kumaran M, Subramanian U, Devarajan B. Performance assessment of variant calling pipelines using human whole exome sequencing and simulated data. BMC Bioinformatics 2019;20:342.

10. Sandmann S, De Graaf AO, Karimi M et al. Evaluating Variant Calling Tools for Non-Matched Next-Generation Sequencing Data. Sci Rep 2017;7:43169.

11. Krøigård AB, Thomassen M, Lænkholm A-V et al. Evaluation of Nine Somatic Variant Callers for Detection of Somatic Mutations in Exome and Targeted Deep Sequencing Data. Jordan IK (ed.). PLOS ONE 2016;11:e0151664.

12. ICGC-TCGA DREAM Somatic Mutation Calling Challenge participants, Ewing AD, Houlahan KE et al. Combining tumor genome simulation with crowdsourcing to benchmark somatic single-nucleotide-variant detection. Nat Methods 2015;12:623–30.

13. Kim SY, Speed TP. Comparing somatic mutation-callers: beyond Venn diagrams. BMC Bioinformatics 2013;14:189.

14. Cai L, Yuan W, Zhang Z et al. In-depth comparison of somatic point mutation callers based on different tumor next-generation sequencing depth data. Sci Rep 2016;6:36540.

15. Xu C. A review of somatic single nucleotide variant calling algorithms for next-generation sequencing data. Comput Struct Biotechnol J 2018;16:15–24.

16. Sandmann S, Karimi M, De Graaf AO et al. appreci8: a pipeline for precise variant calling integrating 8 tools. Hancock J (ed.). Bioinformatics 2018;34:4205–12.

17. Federici G, Soddu S. Variants of uncertain significance in the era of high-throughput genome sequencing: a lesson from breast and ovary cancers. J Exp Clin Cancer Res 2020;39:46.

18. Krøigård AB, Thomassen M, Lænkholm A-V et al. Evaluation of Nine Somatic Variant Callers for Detection of Somatic Mutations in Exome and Targeted Deep Sequencing Data. Jordan IK (ed.). PLOS ONE 2016;11:e0151664.

19. Sandmann S, De Graaf AO, Karimi M et al. Evaluating Variant Calling Tools for Non-Matched Next-Generation Sequencing Data. Sci Rep 2017;7:43169.

20. He X, Chen S, Li R et al. Comprehensive fundamental somatic variant calling and quality management strategies for human cancer genomes. Brief Bioinform 2021;22:bbaa083.

21. Trevarton AJ, Chang JT, Symmans WF. Simple combination of multiple somatic variant callers to increase accuracy. Sci Rep 2023;13:8463.

22. Cantarel BL, Weaver D, McNeill N et al. BAYSIC: a Bayesian method for combining sets of genome variants with improved specificity and sensitivity. BMC Bioinformatics 2014;15:104.

23. Anzar I, Sverchkova A, Stratford R et al. NeoMutate: an ensemble machine learning framework for the prediction of somatic mutations in cancer. BMC Med Genomics 2019;12:63.

24. Ainscough BJ, Barnell EK, Ronning P et al. A deep learning approach to automate refinement of somatic variant calling from cancer sequencing data. Nat Genet 2018;50:1735–43.

25. Majidian S, Agustinho DP, Chin C-S et al. Genomic variant benchmark: if you cannot measure it, you cannot improve it. Genome Biol 2023;24:221.

26. Engle ML, Burks C. Artificially Generated Data Sets for Testing DNA Sequence Assembly Algorithms. Genomics 1993;16:286–8.

27. Alosaimi S, Bandiang A, van Biljon N et al. A broad survey of DNA sequence data simulation tools. Brief Funct Genomics 2020;19:49–59.

28. Stephens ZD, Hudson ME, Mainzer LS et al. Simulating Next-Generation Sequencing Datasets from Empirical Mutation and Sequencing Models. Parkinson J (ed.). PLOS ONE 2016;11:e0167047.

29. Farrell G, Adamidi E, Buono RA et al. Open and Sustainable AI: challenges, opportunities and the road ahead in the life sciences. 2025, DOI: 10.48550/arXiv.2505.16619.

30. Milhaven M, Pfeifer SP. Performance evaluation of six popular short-read simulators. Heredity 2023;130:55–63.

31. Sims D, Sudbery I, Ilott NE et al. Sequencing depth and coverage: key considerations in genomic analyses. Nat Rev Genet 2014;15:121–32.

32. Danecek P, Bonfield JK, Liddle J et al. Twelve years of SAMtools and BCFtools. GigaScience 2021;10:giab008.

33. Cibulskis K, Lawrence MS, Carter SL et al. Sensitive detection of somatic point mutations in impure and heterogeneous cancer samples. Nat Biotechnol 2013;31:213–9.

34. Garrison E, Marth G. Haplotype-based variant detection from short-read sequencing. 2012, DOI: 10.48550/arXiv.1207.3907.

35. Wilm A, Aw PPK, Bertrand D et al. LoFreq: a sequence-quality aware, ultra-sensitive variant caller for uncovering cell-population heterogeneity from high-throughput sequencing datasets. Nucleic Acids Res 2012;40:11189–201.

36. Koboldt DC, Zhang Q, Larson DE et al. VarScan 2: Somatic mutation and copy number alteration discovery in cancer by exome sequencing. Genome Res 2012;22:568–76.

37. Lai Z, Markovets A, Ahdesmaki M et al. VarDict: a novel and versatile variant caller for next-generation sequencing in cancer research. Nucleic Acids Res 2016;44:e108–e108.

38. Kim S, Scheffler K, Halpern AL et al. Strelka2: fast and accurate calling of germline and somatic variants. Nat Methods 2018;15:591–4.

39. Best practises VarDict. https://github.com/AstraZeneca-NGS/VarDict. 2025.

40. Best practises Freebayes. https://github.com/freebayes/freebayes. 2025.

41. Best practises VarScan. https://varscan.sourceforge.net/somatic-calling.html.

42. Best practises LoFreq. https://csb5.github.io/lofreq/commands/.

43. Best practises Mutect2. https://gatk.broadinstitute.org/hc/en-us/articles/360037593851-Mutect2. GATK 2025.

44. Tan A, Abecasis GR, Kang HM. Unified representation of genetic variants. Bioinformatics 2015;31:2202–4.

45. Hasan MS, Wu X, Zhang L. Performance evaluation of indel calling tools using real short-read data. Hum Genomics 2015;9:20.

46. Ghoneim DH, Myers JR, Tuttle E et al. Comparison of insertion/deletion calling algorithms on human next-generation sequencing data. BMC Res Notes 2014;7:864.

47. Guille A, Adélaïde J, Finetti P et al. A benchmarking study of individual somatic variant callers and voting-based ensembles for whole-exome sequencing. Brief Bioinform 2024;26:bbae697.

48. Sisoudiya SD, Tukachinsky H, Keller-Evans RB et al. Tissue-based genomic profiling of 300,000 tumors highlights the detection of variants with low allele fraction. Npj Precis Oncol 2025;9:190.

49. Hollizeck S, Wong SQ, Solomon B et al. Custom workflows to improve joint variant calling from multiple related tumour samples: FreeBayesSomatic and Strelka2Pass. Alkan C (ed.). Bioinformatics 2021;37:3916–9.

50. Jones S, Anagnostou V, Lytle K et al. Personalized genomic analyses for cancer mutation discovery and interpretation. Sci Transl Med 2015;7, DOI: 10.1126/scitranslmed.aaa7161.

51. Roberts ND, Kortschak RD, Parker WT et al. A comparative analysis of algorithms for somatic SNV detection in cancer. Bioinformatics 2013;29:2223–30.

52. Wang M, Luo W, Jones K et al. SomaticCombiner: improving the performance of somatic variant calling based on evaluation tests and a consensus approach. Sci Rep 2020;10:12898.

53. Fragkouli S-C, Iqbal S, Crossman L, et al. An ELIXIR scoping review on domain-specific evaluation metrics for synthetic data in life sciences. 2025, DOI: 10.48550/ARXIV.2506.14508.

